# Comparative analysis of convergent and divergent T7 RNA polymerase promoters for the synthesis of dsRNA *in vivo* and *in vitro*

**DOI:** 10.1101/2025.04.28.650997

**Authors:** Sebastian J. Ross, John Ray, Peter M. Kilby, Mark J. Dickman

## Abstract

Double-stranded RNA plays a key role in various biological processes. The discovery of RNA interference, a gene-silencing mechanism, revolutionised the study of gene function. dsRNA has since been used in novel therapeutics and as an agricultural biocontrol alternative to chemical pesticides. Microbial production typically involves expression systems with convergent T7 promoters. However, convergent transcription from DNA-dependent RNA polymerases can lead to transcriptional interference. In this study, we designed multiple plasmid DNA constructs to investigate the effect of convergent and divergent T7 RNA polymerase production of dsRNA via *in vitro* transcription and *in vivo* in *E. coli*, prior to dsRNA yield quantification and analysis of product quality. We demonstrate that higher yields of larger dsRNA are typically obtained using convergent promoters during *in vivo* production. A typical fold increase of 2.1 was obtained for dsRNA > 400 bp. However, production of smaller dsRNA (< 250 bp) by divergent promoters resulted in increased yields (2.2 fold). Furthermore, our data demonstrates that *in vitro* transcription production of dsRNA using divergent T7 promoters results in significantly higher yields of dsRNA, with a maximum fold increase of 6.46. Finally, independent of size, we demonstrate that dsRNA synthesised from DNA templates with multiple transcriptional terminators, improved the quality and purity of dsRNA due to decreased formation of dsRNA multimers or aggregates, compared to run-off transcription. This study highlights alternative optimal strategies for the production of a wide range of different sized dsRNA both *in vitro* and in microbial systems.

## Introduction

Double-stranded ribonucleic acid (dsRNA) is a ubiquitous molecule found throughout nature. It plays a fundamental role in several crucial biological processes, including gene regulation and stability, and antiviral defence mechanisms, via the mechanism of RNA interference (RNAi) (Elliot and Ladomery 2016; Mamta and Rajam 2017; Chen and Hur 2022). In recent years dsRNAs, have revolutionised the understanding and study of gene function. Of particular significance is their capacity to initiate RNAi pathways, conserved across eukaryotes, facilitating precise gene silencing and suppression, and presenting innovative strategies for studying and addressing human diseases and managing plant pests (Baum et al. 2007; Mao et al. 2007; Davis et al. 2010; Adams et al. 2018; Nguyen et al. 2018; Lopez et al. 2019).

The mechanism of gene silencing or suppression via antisense RNA was observed in early gene expression studies (Mol et al. 1990), and plant pigmentation studies, coined co-suppression (Napoli et al. 1990). Fire et al. (1998) were the first to characterise RNAi during studies on *Caenorhabditis* elegans, in which the addition of sequence-specific long double-stranded RNA (dsRNA) led to an observed phenotypic change as the result of messenger RNA (mRNA) degradation, a form of post-transcriptional gene silencing. In 2001, short dsRNAs or short interfering RNA (siRNA), 21-24 nucleotides, were demonstrated to be effective in silencing within mammalian cells without inducing an interferon response (Elbashir et al. 2001).

The utilization of dsRNA as a therapeutic in an animal model was first demonstrated by McCaffrey et al. (2002) via the inhibition of hepatitis C virus replication in mice, using siRNA and small hairpin dsRNA (hpRNA). This led to a proliferation of investigations into the therapeutic applications of dsRNA. By 2010 the first RNAi clinical trials took place, investigating a treatment for widespread melanoma, by targeting the M2 subunit of a ribonucleotide reductase (Davis et al. 2010). Davis et al. (2010), used a nanoparticle encapsulated dsRNA, demonstrating successful cleavage of the target mRNA. In 2018, the first siRNA drug, Patisiran, was approved for the treatment of hereditary transthyretin amyloidosis following successful stage 3 clinical studies (Adams et al. 2018).

Within the agricultural industry, RNAi has been demonstrated as a successful crop protection system. Initial studies demonstrated successful gene expression interference via both applications of synthetically produced dsRNA, as well as endogenous dsRNA produced through transgenic plants encoding dsRNA insecticidal genes (Dong and Friedrich 2005; Baum et al. 2007; Mao et al. 2007). These proof-of-concept studies highlighted the potential of dsRNA as a crop protection system via transgenic methods but also via the application of dsRNA as a biopesticide. Since then, scientists have demonstrated successful RNAi in numerous insect pests across a range of orders and species, through several different application methods (Li et al. 2013; San Miguel and Scott 2016; Vyas et al. 2017; Christiaens et al. 2018; Mogilicherla et al. 2018; Nguyen et al. 2018; Lopez et al. 2019). More recently, as of January 2024, GreenLight Biosciences received approval from the U.S. Environmental Protection Agency (EPA) for the first commercially available dsRNA RNAi biocontrol, Calantha^TM^ (https://www.calanthaag.com).

Production of long dsRNA, siRNA, and hpRNA, varies depending on the application. RNAs are traditionally produced either via run-off *in vitro* transcription (IVT), *in vivo* within microbial cells, endogenously within transgenic plants, or by cell-free expression, using plasmid DNA or PCR products as templates and T3, T7, or Sp6 RNA polymerases (RNAP) (Timmons et al. 2001; Rajagopal et al. 2002; Alder et al. 2003; Skelly et al. 2003; Maxwell et al. 2018; Wang et al. 2018; Howard et al. 2022). *In vitro* production provides the added benefit of easily incorporating a wide range of chemically modified bases to increase the efficiency of RNAi due to a decrease in dsRNA degradation caused by target organism nucleases (Braasch et al. 2003; Soutschek et al. 2004; Peacock et al. 2011; Lima et al. 2012; Howard et al. 2022).

Various designs have been used to produce dsRNA, including the original dsRNA plasmid, L4440, which consisted of two convergent T7 promoters flanking a gene of interest (GOI) (Timmons et al. 2001), DNA templates including a single T7 promoter to produce hpRNA (Dalakouras et al. 2018), or the two individual templates to produce the sense and antisense strands, which are annealed in equal molar quantities post purification (Howard et al. 2022). More recent plasmid designs for *in vivo* production of long dsRNAs and hpRNAs have incorporated transcriptional terminators, demonstrating increased yields of up to a 2-fold increase in hpRNA (Chen et al. 2019), and a 7.8-fold increase of dsRNA when utilising three consecutive terminators (Ross et al. 2024).

The conventional approach for producing dsRNA *in vivo,* involves the use of two convergent T7 promoters flanking a target sequence. Convergent transcription is thought to play a significant role in gene regulation within organisms, found typically within extra-chromosomal elements (Callen et al. 2004). However, opposing convergent promoters can lead to the reduction of transcription in prokaryotic and eukaryotic systems. (Ward and Murray 1979; Horowitz and Platt 1982; Elledge and Davis 1989; Eszterhas et al. 2002; Lehner et al. 2002; Prescott and Proudfoot 2002; Callen et al. 2004).

The effect of transcriptional interference has been studied using *E. coli* RNAP and T7 RNAP. Callen et al. (2004), observed a 5.6-fold interference between convergent strong and weak promoters and a 1-fold interference between divergent promoters when using the multi-subunit *E. coli* RNAP holoenzyme. Furthermore, terminating transcriptional prior to opposing promoter region reduced transcriptional interference, indicating a mechanism of interference generated by translocation of RNAPs and their opposing promoter regions (Callen et al. 2004). A “sitting duck” mechanism, which involves head-on collisions of elongating RNAP from one strand and initiation intermediates from the opposite promoter was proposed (Callen et al. 2004). However, it is argued that this would not greatly inhibit transcription as a new RNAP would rapidly bind (Sneppen et al. 2005).

Crampton et al. (2006) investigated three mechanisms of transcriptional interference: promoter occlusion (previously proposed by Sherwin et al. (2005)), “sitting duck,” and elongation complex (EC) collisions within target DNA. Using *E. coli* RNAP-σ70, Crampton et al. (2006) observed that RNAP collisions often resulted in stalled RNAPs, sometimes leading to backtracking and transcriptional roadblocks. EC collisions can also cause RNAP detachment, producing truncated RNA transcripts (Chatterjee et al. 2011). Hobson et al. (2012) found that convergent RNAPII ECs are unable to pass each other during transcription following collison and remain bound to DNA *in vitro.* However, they are removed *in vivo* within yeast through ubiquitylation-directed proteolysis.

Wang et al. (2023) developed a single-molecule assay to study *E. coli* RNAP on DNA, showing that head-on collisions prevent read-through, but do not cause RNAP release. However, co-directional collisions into stalled convergent ECs significantly enhance termination efficiency. Additionally, stem-loop structures in nascent RNA help localize collisions between convergent genes, suggesting that RNAP collisions contribute to transcriptional termination and gene regulation (Wang et al. 2023).

Previous studies of the single sub-unit RNAPs, T7 and SP6, demonstrated their ability to displace DNA-bound proteins during transcription (Campbell and Setzer 1992; Clark and Felsenfeld 1992; Gallego et al. 1995). Zhou and Martin (2006), investigating co-directional T7 RNAP, demonstrated that stalled T7 RNAPs are displaced by trailing RNAPs. Furthermore, T7 RNAP binding is prevented if the leading complex is 12 bp or less from the promoter. Displacement of the leading complex is only possible at 20 bp from the promoter. In addition, it was shown that following a convergent collision of two single subunit RNAPs T7 and T3, the RNAPs can incrementally pass each other and largely retain transcriptional activity. Finally, their results suggest the RNAPs temporarily release the non-template strand (Ma and McAllister 2009).

In this study, we designed a range of novel plasmid DNA constructs to study the effect of convergent and divergent T7 RNAP promoters on the production of dsRNA both in microbial cells and via IVT. This allowed us to investigate the effect of the orientation and spatial location of the T7 promoters on the total dsRNA yield and quality of dsRNA.

## Results and Discussion

### Investigating the effect of convergent and divergent T7 RNAP promoters on the production of dsRNA biocontrols in *E. coli*

### Plasmid construct design

Multiple plasmids were used to investigate the difference in production of dsRNA in *E. coli* HT115(DE3), between convergent and divergent T7 RNAP promoters. The convergent promoter systems consist of a single dsRNA gene sequence flanked by two T7 RNAP promoters, finally flanked by a combination of three transcriptional terminators, T7 S + *rrnB* T1 + T7 S (3T) (Mairhofer et al. 2015; Calvopina-Chavez et al. 2022; Ross et al. 2024). The convergent promoter systems were developed and previously utilised by Ross et al. (2024). The divergent promoter systems utilise the same dsRNA sequences, flanked by a single T7 RNAP promoter on one side and three transcriptional terminators on the other, separated by a spacer region and orientated in opposite directions to avoid polymerase crossover (see Figure 1).

**Figure 1.**
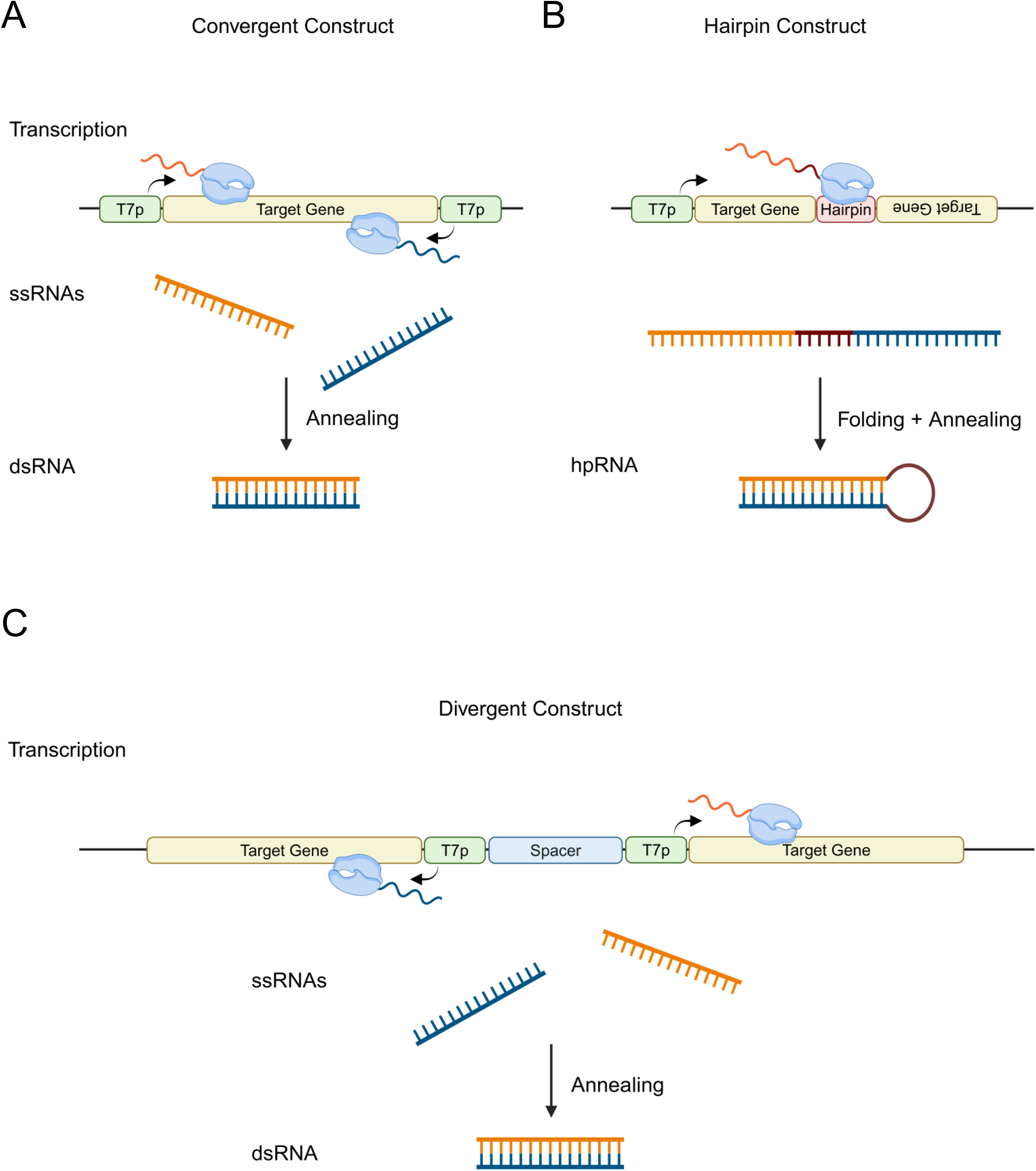
Schematic illustration of DNA templates for the synthesis of dsRNA. Schematics representing three constructs and methods for the synthesis of dsRNA. These include convergent, hairpin and divergent production. **A.** The convergent construct consists of a target gene flanked by two T7 polymerase promoters orientated in opposite directions, leading to simultaneous transcription of both sense and antisense ssRNA of the target sequence, via convergent transcription. ssRNAs then anneal post transcription leading to the formation of the dsRNA. **B.** The hairpin construct consists of a single T7 polymerase promoter upstream of a target gene. Downstream of the target gene a hairpin sequence is located followed by the target gene in a reverse orientation. A single ssRNA is produced which subsequently folds and anneals to its complementary base pairs, forming the hpRNA. **C.** The divergent construct is formed on a target gene flanked by a T7 polymerase promoter downstream orientated in the antisense direction. Next a spacer region of X bp separates the first T7 polymerase promoter from another T7 polymerase promoter, which is located upstream of a second target gene sequence, orientated in the sense direction. Two ssRNAs are transcribed simultaneously by two diverging T7 polymerase promoters. Post transcription the ssRNAs anneal to form the target dsRNA. Created in BioRender. Dickman, M. (2025) https://BioRender.com/j93g547

Initial work focused on the drosophila dsRNA sequence Dome 11 (400 bp) to generate the convergent promoter plasmid (pD11_C) and the divergent promoter plasmid with a spacer region of 225 bp (pD11_D). In addition, plasmid constructs were also designed to vary the spacer region between the two divergent T7 RNAP promoters (pD11_D_1kb) and (pD11_D_2kb), by incorporating random sequences into the spacer region of pD11_D. All plasmids utilised the vector pMA-7, containing a ColEI origin of replication. A schematic of plasmid constructs used in this study can be found in Supplementary Figure S1.

### Increased yield of Dome11 dsRNA using convergent T7 RNAP promoters in *E. coli*

Dome11 plasmids (pD11_C, pD11_D, pD11_D_1kb and pD11_D_2kb), were transformed into *E. coli* (HT115) cells, prior to cell growth and induction with IPTG. The corresponding growth curves are shown in Figure 2A and show similar growth curves for both the convergent and divergent constructs, with final mean OD_600_ values ranging from 1.33 to 1.36 (Figure 2A). RNA extractions were performed on 1 x 10^9^ cells, calculated from the final OD_600_. RNA extractions and purifications were performed in both the presence and absence of RNase T1 to remove endogenous *E. coli* ssRNA (rRNA and tRNA) prior to agarose gel electrophoresis analysis (Figure 2B).

**Figure 2.**
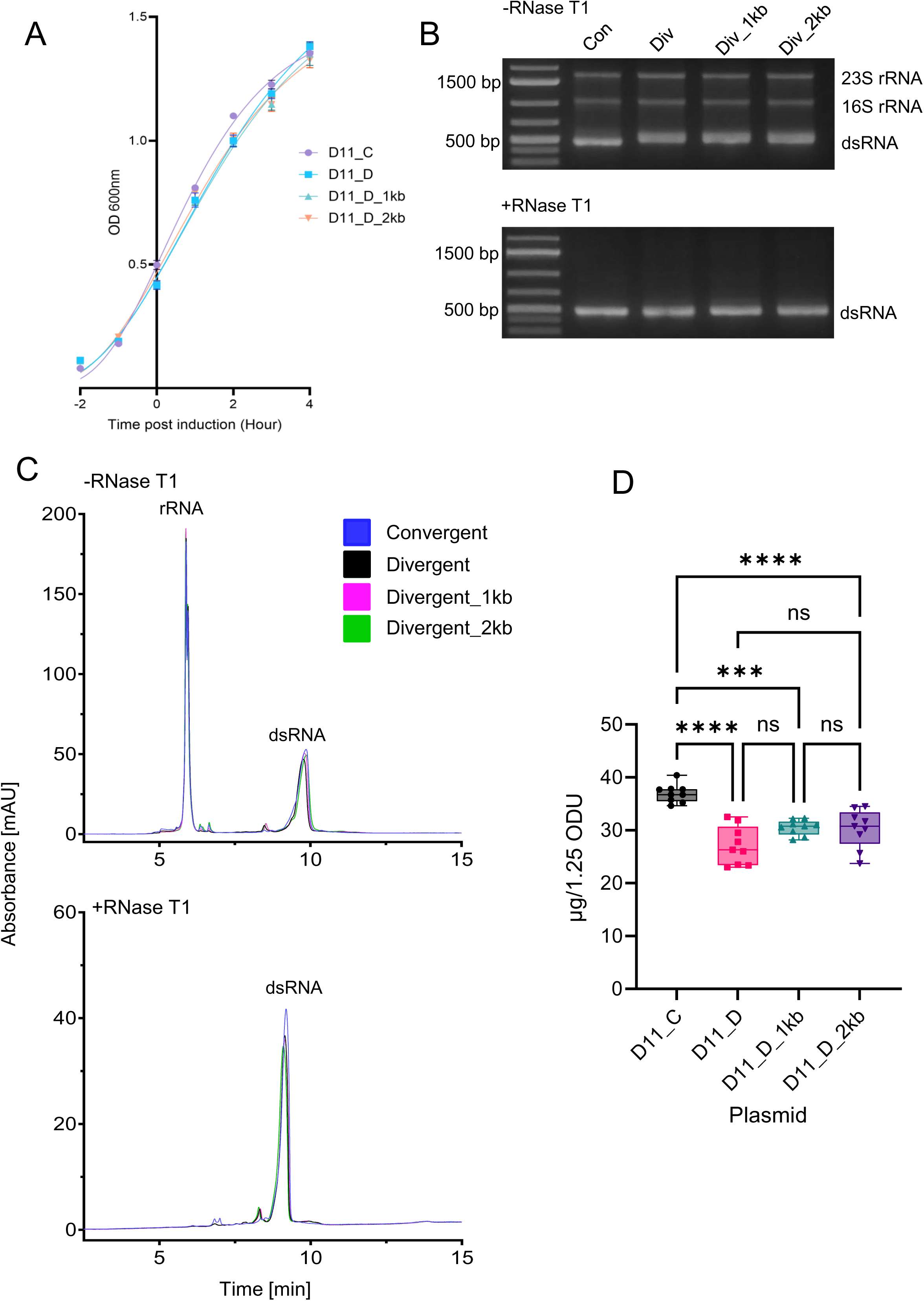
Investigations of convergent and divergent synthesis of Dome11 dsRNA *in vivo* within *E. coli*. Plasmids D11_C, D11_D, D11_D_1kb and D11_D_2kb were investigated **A.** Growth curve comparison of *E. coli* HT115 cells transformed with each of the plasmids. An outgrowth was performed and allowed to grow to an OD_600_ of 0.4-0.6. Samples were then induced for 4 hours, with OD measurements recorded at hour time points. Data shown as mean ± SD and are representative of three biological replicates. **B.** Agarose gel electrophoresis analysis of RNA extractions in the absence and presence of RNase T1. The corresponding rRNA and Dome 11 dsRNA (400 bp) are highlighted. **C.** IP-RP HPLC chromatography analysis in the absence and presence of RNase T1. **D**. Quantification of dsRNA yield. RNA extractions were performed in the presence of RNase T1. RNA samples were analysed using UV spectrophotometry to determine RNA concentration using a mass concentration/A260 unit (46.52 μg/mL). Significance was calculated using a one-way ANOVA test with multiple comparisons. Convergent-produced Dome11 (pD11_C) demonstrates the highest absolute yield of 36.95 µg. Data is shown as box plots of triplicate technical replicates and are representative of three biological replicates; p<0.05 = (*), p<0.01 = (**), p<0.001 = (***), p<0.0001 (****).

The results show the successful production of dsRNA Dome11 (≈400 bp) by each of the plasmid designs. Semi-quantitative analysis based on gel band intensity on extracts both in the absence and presence of RNase T1 indicates that production using convergent promoters (pD11_C), results in a small increase in dsRNA yield. Differences in band migration are noted between the two constructs in the absence of RNase T1, which is removed in the presence of RNase T1. These results indicate that the difference is due to ssRNA overhangs present on the dsRNA generated using the divergent promoters potentially due to a decrease in termination efficiency resulting in longer ssRNA overhangs (Figure 2B).

Accurate quantification of dsRNA yield was performed via a combination of relative quantification of dsRNA (IP-RP-HPLC analysis) and absolute quantification (UV spectrophotometry), following purification of dsRNA (Ross et al. 2024). IP-RP-HPLC analysis of the RNA extracted is shown in Figure 2C and includes both the dsRNA and *E. coli* rRNA from convergent and divergent promoter constructs.

Absolute quantification of the Dome11 dsRNA was performed following extraction of dsRNA in the presence of RNase T1 to remove the ssRNA prior to UV spectrophotometry analysis (Ross et al. 2024) (see Figure 2D). Total dsRNA analysis demonstrated mean total dsRNA yields of 36.95 µg, 27.10 µg, 30.50 µg and 30.29 µg (from 1 x 10^9^ cells), between the convergent (pD11_C) and divergent (pD11_D, pD11_D_1kb and pD11_D_2kb) systems, respectively. The results show a fold decrease of 0.73, 0.83 and 0.82, respectively. These results are consistent with relative quantification (see Supplementary Figure S2).

These results show that utilising convergent T7 RNAP promoters results in an increase in the dsRNA yield from *E. coli* HT115 cells compared to divergent T7 RNAP promoters. Furthermore, we have demonstrated that the size of the spacer region between the two T7 RNAP promoters in the divergent promoter system does not have a significant effect on the production of dsRNA.

### Optimisation of dsRNA yield using convergent and divergent T7 RNAP promoters is dependent on the size of dsRNA

Further studies were performed to investigate the effect of the *in vivo* production of alternative dsRNA gene sequences, GFP (263 bp) and Sec23A (1504 bp), using either a convergent or divergent promoter system. These alternative smaller and larger sequences were chosen to investigate the effect of alternative dsRNA sequences on the dsRNA yield. Furthermore, Sec23A was chosen to investigate the effect of a divergent system on dsRNA quality, as previous unpublished data demonstrated that the production of Sec23 dsRNA resulted in a range of dsRNA length impurities.

The plasmids were transformed into *E. coli* (HT115) cells, prior to cell growth and induction with IPTG. The corresponding growth curves are shown in Figure 3A. The GFP dsRNA growth curve data indicates a small increase in metabolic burden on the cells when utilising a divergent production, with a final mean OD_600_ of 1.33 compared to 1.53 in the convergent system. Sec23A indicates similar metabolic differences with final mean OD_600_ measurements of, 1.27 and 1.24 for between convergent and divergent constructs, respectively.

**Figure 3.**
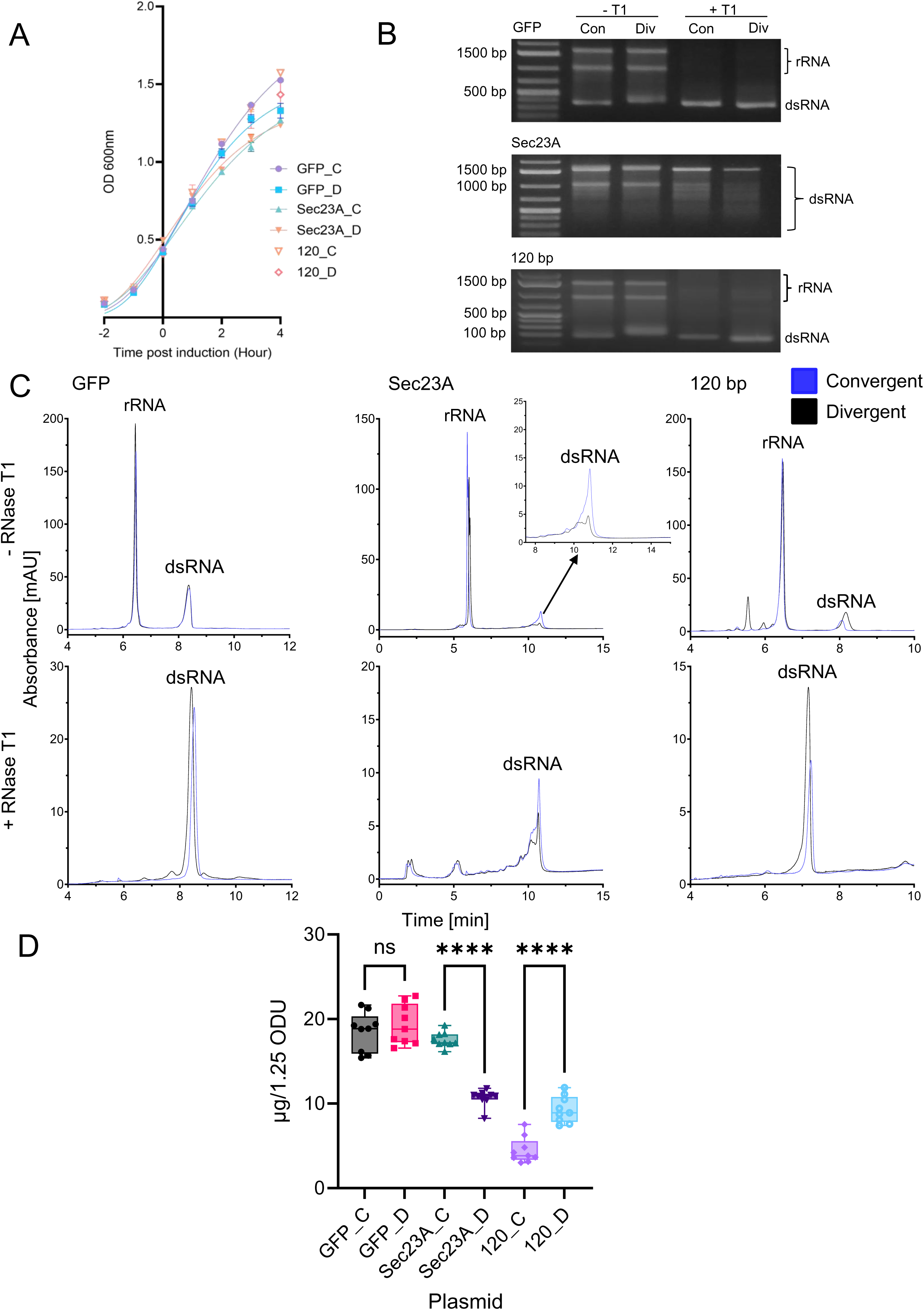
Investigations of convergent and divergent synthesis of GFP, Sec23A and 120 bp dsRNA *in vivo* within *E. coli*. Plasmids GFP_C, GFP_D, Sec23A_C, Sec23A_D, 120_C and 120_D were investigated **A.** Growth curve comparison of *E. coli* HT115 cells transformed with each of the plasmids. An outgrowth was performed and allowed to grow to an OD_600_ of 0.4-0.6. Samples were then induced for 4 hours, with OD measurements recorded at hour time points. Data shown as mean ± SD and are representative of three biological replicates. **B.** Agarose gel electrophoresis analysis of RNA extractions in the absence and presence of RNase T1. The corresponding rRNA and GFP (263 bp), Sec23A (1504 bp) and 120 bp dsRNA are highlighted. **C.** IP-RP HPLC chromatography analysis in the absence and presence of RNase T1. Insert is shown for Sec23A dsRNA in the absence of RNase T1. **D**. Quantification of dsRNA yield. RNA extractions were performed in the presence of RNase T1. RNA samples were analysed using UV spectrophotometry to determine RNA concentration using a mass concentration/A260 unit (46.52 μg/mL). Significance was calculated using unpaired T-tests, against each gene sequence pairing. No significant difference was noted between GFP constructs. Convergent-produced Sec23A (pSec23A_C) demonstrated the highest absolute yield (17.53 µg) between Sec23A constructs. Divergent-produced 120 bp (p120_D) demonstrated the highest absolute yield (9.29 µg) between 120 bp constructs. Data is shown as box plots of triplicate technical replicates and are representative of three biological replicates; p<0.05 = (*), p<0.01 = (**), p<0.001 = (***), p<0.0001 (****).

Agarose gel electrophoresis of dsRNA extracted for GFP and Sec23A (263 bp and 1504 bp, respectively) in the presence and absence of RNase T1 are shown in Figure 3B. Interestingly for the smaller dsRNA (GFP), a higher migration band is noted for the divergent plasmid construct in the absence of RNase T1 consistent with the data from Dome11 dsRNA (see Figure 2B). The results for Sec23A dsRNA demonstrate the presence of various smaller bands that are visible in the absence and presence of RNase T1, this indicates that these bands are shorter dsRNAs, which are potentially due to early termination or degradation of ssRNA from partially annealed Sec23A dsRNA.

Semi-quantitative analysis based on gel band intensity for GFP indicates similar levels of dsRNA production from both convergent and divergent promoters. Sec23A data indicates an increase in dsRNA yield of both the full-length dsRNA and shorter dsRNAs produced from the convergent promoter system. Quantitative analysis of the main full-length Sec23A dsRNA was performed using densitometry analysis and shows a 2.22-fold difference in full-length Sec23A dsRNA.

Absolute quantification demonstrated no significant difference in mean total dsRNA yields, 18.49 µg and 19.34 respectively between pGFP_C and pGFP_D (Figure 3D). Comparison between mean total dsRNA yields of pSec23A_C (17.53 µg) and pSec23A_D (10.69 µg) demonstrated a 0.61-fold decrease in mean total dsRNA yields when produced within the divergent system, consistent with the relative quantification (Supplementary Figure S3).

Based on our previous *in vivo* findings where we demonstrated a significant increase in yield when producing larger dsRNAs (Dome11 and Sec23A) and no significant difference in yield for the smaller dsRNA (GFP) when using a convergent system, we decided to investigate a smaller 120 bp dsRNA sequence. We hypothesised that potentially an even smaller sequence would lead to an increase in dsRNA yield when utilising a divergent T7 RNAP promoters.

Two new constructs, p120_C and p120_D, were produced and transformed into *E. coli* (HT115) cells, prior to cell growth and induction with IPTG. The corresponding growth curves are shown in Figure 3A. The growth curve patterns were similar to those seen with the smaller GFP constructs, pGFP_C and pGFP_D, in which p120_C indicated a higher final mean OD_600_ of 1.57, compared to p120_D, 1.43.

Agarose gel electrophoresis analysis of the extracted 120 bp dsRNA from both constructs, in the presence and absence of RNase T1 are shown in Figure 3B. Semi-quantitative analysis based on gel band intensity indicated a small increase in dsRNA yield for divergent promoter constructs compared to p120_C and higher migrating bands. The increased band intensity is a novel result for *in vivo* production using a divergent T7 system, while the higher band migration noted in the absence of RNase T1 is comparable to previous results from divergent constructs, pD11_D, pD11_1kb, pD11_2kb and pGFP_D.

Further quantitative analysis via relative and absolute quantification of dsRNA was performed as previously described. IP-RP chromatograms of the extracted RNA in the absence of RNase T1 indicate differences in product sizes between the p120_C and p120_D produced dsRNA (Figure 3C). The increased product size of p120_D dsRNA could potentially be due to additional single-stranded overhangs. This is proposed as following analysis of the IP-RP-HPLC traces of RNase T1 treated dsRNA, the peaks align more closely. There is a slight increase in retention time for p120_C samples, but this can be explained by the size of the overhangs being 6 bp larger following RNase T1 digestion due to the final guanine location. Absolute mean dsRNA yields between p120_C (4.46 µg) and p120_D (9.29 µg), were consistent with the results of the relative quantification (Supplementary Figure S3), demonstrating a fold increase of 2.08 when using the divergent T7 promoter system (See Figure 3D).

In summary, the results show that an increase in yield of the larger dsRNA, Sec23A, was observed using convergent T7 promoter systems in *E. coli* HT115(DE3) cells, consistent with Dome11. The results showed typically a 1.3 – 2.1-fold increase in relative and absolute abundance of dsRNA for two different dsRNA sequences used in this study. However, no significant increase in yield was obtained for the smaller GFP dsRNA sequence when comparing different T7 promoter systems. Furthermore, we have demonstrated that *in vivo* production of a small 120 bp dsRNA is significantly increased when using divergent T7 promoters.

These results indicate that the size of the dsRNA plays a key role deciding which promoter system is optimal for the synthesis of dsRNA. It is proposed that for the production of smaller dsRNA sequences a higher rate of T7 RNAP collisions may occur during convergent T7 RNAP production due to the ratio of RNAP per bp of DNA template. This leads to increased transcriptional interference via termination of transcriptions and blocking of polymerase-promoter binding (Callen et al. 2004; Shearwin et al. 2005; Chatterjee et al. 2011; Wang et al. 2023).

However, these results are in contrast to a previous *in vivo* study which increasing the space between strong and weak convergent promoters by 102 bp led to an increase in transcriptional interference (Callen et al. 2004). It should be noted that these studies used the multisubunit *E. coli* RNAP holoenzyme, not T7 RNAP, and unequal strength promoters in some cases. Furthermore, a study using the bacteriophage RNAPs, T7 and T3, suggests that bacteriophages can incrementally walk past each other following collision (Ma and McAllister 2009). However, this was performed *in vitro* and is therefore likely to perform differently *in vivo*.

Finally, it is proposed that more efficient annealing of larger ssRNAs produced using convergent T7 promoters, compared to divergent T7 RNAP promoters, is due to their spatial proximity within the cell. This enables formation of the corresponding dsRNA before degradation of the ssRNA by endogenous RNases. Therefore, it is suggested that more favourable annealing of the ssRNA due to spatial proximity outweighs the effect of transcriptional interference of the T7 RNAPs resulting in increased yield of larger dsRNAs via convergent T7 RNAP promoters. In contrast, optimal production of smaller dsRNA occurs using divergent T7 RNAP promoters.

### Investigating the effect of convergent and divergent T7 RNAP systems on *in vitro* transcription of dsRNA

In addition to the production of dsRNA biocontrols in microbial cells, an alternative method for their production is via enzymatic synthesis using *in vitro* transcription (IVT).These are largely performed for smaller-scale studies (Timmons et al. 2001; Wang et al. 2018; Howard et al. 2022), but can also be used in large-scale manufacturing of dsRNA (Genolution, South Korea, https://genolution.co.kr/). For IVT production of dsRNA, templates can include plasmid or PCR templates using convergent T7 RNAPs, a single T7 RNAP hairpin, or two individual templates to produce both ssRNAs, which are annealed post-production (Timmons et al. 2001; Dalakouras et al. 2018; Hough et al. 2022; Howard et al. 2022).

To investigate the effect of convergent and divergent T7 RNAP promoters for the *in vitro* production of dsRNA, plasmid DNA templates previously described were linearised with a restriction enzyme that produced a cut in the ampicillin marker in the plasmid backbone, producing linear templates containing the transcriptional terminators (see Supplementary Figure S4). Alternatively, restriction digestion was performed to remove transcriptional terminators and generate linearised templates at the end of the dsRNA gene sequence for run-off *in vitro* transcription (see Supplementary Figure S4).

### Increased production of dsRNA in vitro using divergent promoters in conjunction with multiple transcriptional terminators

IVT reactions were performed to produce dsRNA from linearised DNA templates for each of the convergent and divergent promoter systems, for genes Dome11, GFP, Sec23A, and the 120 bp dsRNA sequence. In each case, multiple transcriptional terminators were present at the end of the dsRNA gene sequences similar to those plasmids previously used in the production of dsRNA in *E. coli*. IVT reactions were performed using equimolar amounts of DNA template (250 fmol) and the samples were digested with RNase T1 to remove ssRNA overhangs prior to downstream purification and analysis. Agarose gel electrophoresis shows the successful production of dsRNA for all four of the target sequences using both divergent and convergent promoters (see Figure 4A). Semi-quantitative analysis based on gel band intensity indicated increased dsRNA yields for all divergent T7 RNAP promoter systems, compared to the convergent. IVT of Sec23A resulted in the production of a full-length dsRNA target sequence, along with various smaller dsRNAs consistent with previous *in vivo* production. Densitometric analysis of the main full-length Sec23A dsRNA indicated a 2.56- fold difference in full-length Sec23A dsRNA.

**Figure 4.**
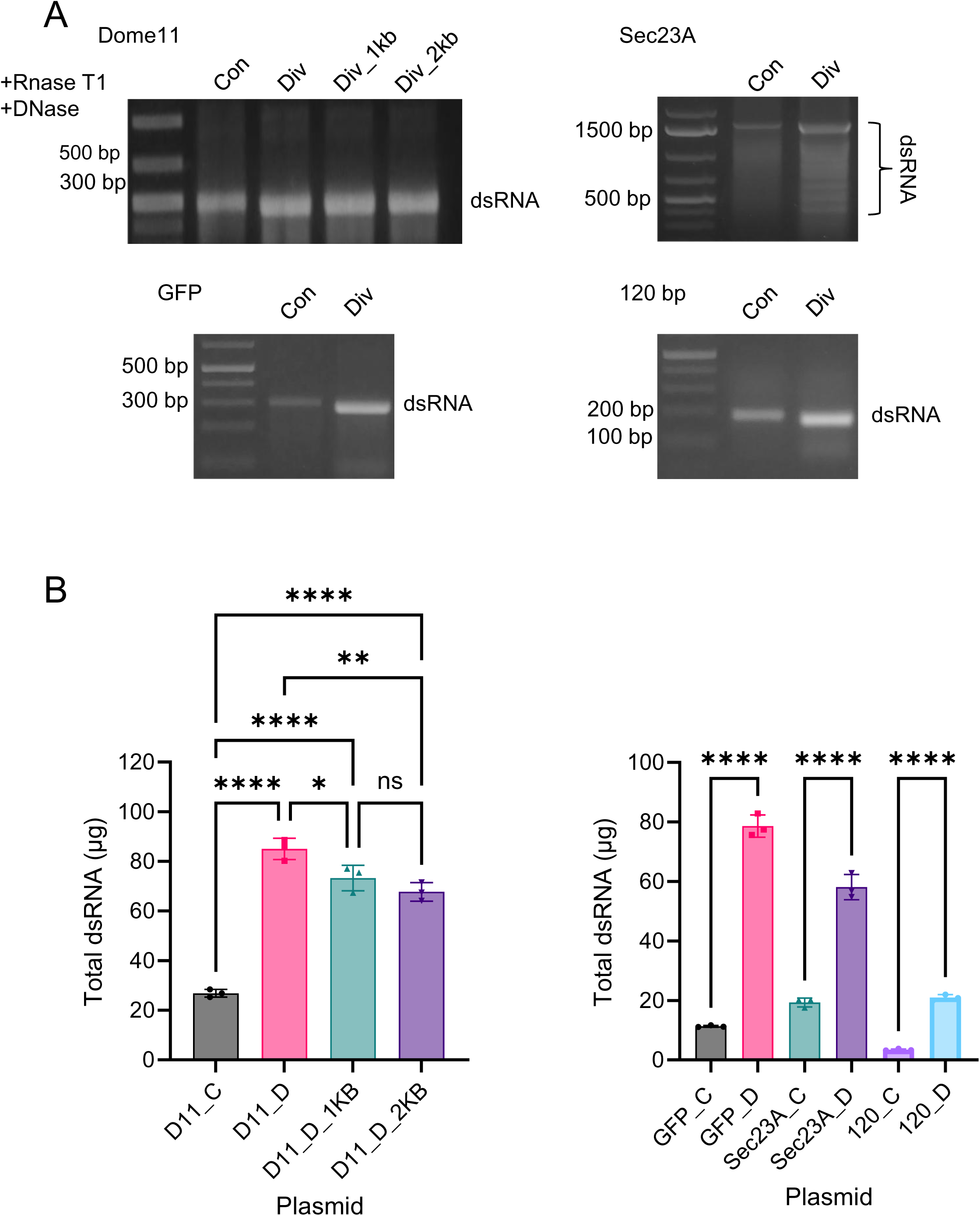
Investigations of convergent and divergent synthesis of Dome11, GFP, Sec23A and 120 bp dsRNA *in vitro* utilising transcriptional terminators. All Dome11, GFP, Sec23A and 120 bp plasmids constructs were investigated **A.** Agarose gel electrophoresis analysis of RNA extractions in the presence of RNase T1 and DNase. The corresponding Dome 11 (400 bp), GFP (263 bp), Sec23A (1504 bp) and 120 bp dsRNA are highlighted. **B**. Quantification of dsRNA yield. RNA samples were generated from 20 µl IVT reactions, followed by the addition of RNase T1 and DNase prior to purification. RNA samples were analysed using UV spectrophotometry to determine RNA concentration using a mass concentration/A260 unit (46.52 μg/mL). Significance was calculated using a one-way ANOVA test with multiple comparisons, for Dome11 RNA samples. Divergent-produced Dome11 (pD11_D) demonstrates the highest absolute yield of 84.99 μg. Significance was calculated using unpaired T tests, for GFP, Sec23A and 120 bp RNA samples. Divergent-produced, GFP (pGFP_D), Sec23A (pSec23A_D), and 120 bp (p120_D) demonstrate the highest absolute yields of, 78.64 μg, 58.12 μg, and 20.95 μg, respectively. Data is shown as bar charts of triplicate technical replicates; p<0.05 = (*), p<0.01 = (**), p<0.001 = (***), p<0.0001 (****).

Quantitative analysis via relative and absolute quantification of dsRNA was performed as previously described and summarised in Figure 4B and Supplementary Figure S5/6. Absolute quantification of dsRNA (total dsRNA mass from a 20 μl IVT reaction) was consistent with the relative quantification analysis. A significant increase in mean relative Dome11 dsRNA yields was observed utilising divergent T7 RNAP promoters with the greatest mean fold increase of 3.16 obtained. However, a small decrease in dsRNA yield was observed in the divergent promoter system as the spacer length increased from 225-2000 bp. Similarly, a significant increase in mean absolute dsRNA yield was recorded for GFP, Sec23A and 120bp. Mean fold increases of 6.96, 3.01 and 6.14 were recorded respectively. These results are consistent with relative quantification (see Supplementary Figure S6)

In summary, we have demonstrated that increased yields of dsRNA are produced via IVT reactions from DNA templates utilising multiple transcriptional terminators using divergent T7 RNAP promoters compared to the convergent promoters. These findings are consistent with increased transcriptional interference of T7 RNAP that occurs using convergent promoters compared to divergent promoters (Callen et al. 2004; Crampton et al. 2006; Hobson et al. 2012; Wang et al. 2023). Finally, we have demonstrated that the spacer region between divergent promoters negatively affects the production of *in vitro* transcribed dsRNA when the region is over 1 kb in size when utilising transcriptional terminators for termination in IVT reactions.

### Increased production of dsRNA from divergent promoters using run-off *in vitro* transcription

Further analysis of convergent and divergent promoters on the IVT of dsRNA was performed using alternative DNA templates linearised after the target gene sequence in the absence of multiple transcriptional terminators. IVT reactions were performed as previously described for genes Dome11, GFP, Sec23A and the 120 bp sequence using run-off transcription. Samples were digested with RNase T1 prior to analysis using gel electrophoresis and IP-RP-HPLC (See Figure 5A and Supplementary Figure S7). Semi-quantitative analysis based on gel band intensity indicated higher dsRNA yields for all divergent promoter systems. In addition, Dome11, GFP and 120 bp dsRNA show the presence of additional bands corresponding to larger dsRNA species in both the agarose gel electrophoresis (above the main dsRNA, see Figure 5A) and also in the IP-RP HPLC (eluting later than the main dsRNA, see Supplementary Figure S7). This is potentially due to the formation of dsRNA multimers or aggregates as previously observed (Nwokeoji et al. 2019). The analysis of Sec23A dsRNA shows the production of the full-length dsRNA target sequence and various smaller bands, as previously demonstrated (see Figure 4A). Densitometric analysis of the main full-length Sec23A dsRNA indicated a 4.25-fold increase in full-length Sec23A dsRNA using the divergent T7 RNAP system.

**Figure 5.**
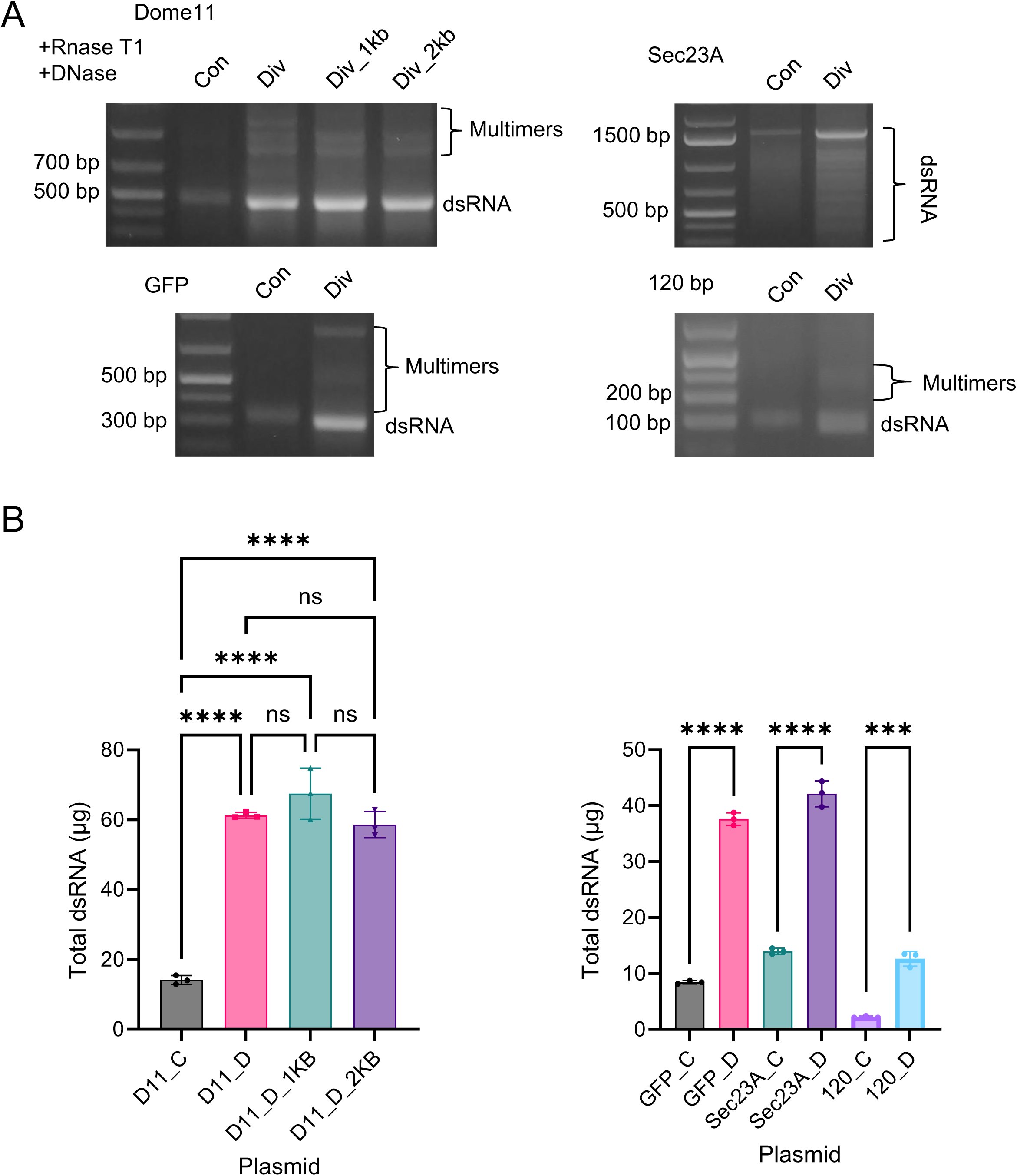
Investigations of convergent and divergent synthesis of Dome11, GFP, Sec23A and 120 bp dsRNA *in vitro* utilising run-off transcription. All Dome11, GFP, Sec23A and 120 bp plasmids constructs were investigated **A.** Agarose gel electrophoresis analysis of RNA extractions in the presence of RNase T1 and DNase. The corresponding Dome 11 (400 bp), GFP (263 bp), Sec23A (1504 bp) and 120 bp dsRNA and potential multimers are highlighted. **B**. Quantification of dsRNA yield. RNA samples were generated from 20 µl IVT reactions, followed by the addition of RNase T1 and DNase prior to purification. RNA samples were analysed using UV spectrophotometry to determine RNA concentration using a mass concentration/A260 unit (46.52 μg/mL). Significance was calculated using a one-way ANOVA test with multiple comparisons for Dome11 RNA samples. Divergent-produced Dome11 (pD11_D_1kb) demonstrates the highest absolute yield of 67.44 μg. Significance was calculated using unpaired T tests, for GFP, Sec23A and 120 bp RNA samples. Divergent-produced, GFP (pGFP_D), Sec23A (pSec23A_D), and 120 bp (p120_D) demonstrate the highest absolute yields of, 37.61 μg, 42.13 μg, and 12.63 μg, respectively. Data is shown as bar charts of triplicate technical replicates; p<0.05 = (*), p<0.01 = (**), p<0.001 = (***), p<0.0001 (****).

Quantitative analysis via relative and absolute quantification of dsRNA was performed as previously described and are summarised in Figure 5B and Supplementary Figure S8. Dome11 dsRNA demonstrated the greatest significant mean fold increase of 4.77, between pD11_C and pD11_D_1kb. GFP, Sec23A and 120 bp dsRNA demonstrated significant mean fold increases of, 4.46, 3.01 and 5.66, respectively, from divergent promoters in comparison to convergent promoters. These results are consistent with relative quantification (see Supplementary Figure S8).

Finally, several studies were performed to investigate the production of chemically modified dsRNA by the incorporation of N1-methylpseudouridine as a replacement for uridine, via either convergent or divergent promoters (See Supplementary Figure S9/10). The results show an increase in chemically modified dsRNA yields using divergent promoters consistent with previous results.

In summary, we have demonstrated that yields of dsRNA produced in IVT reactions utilising either run-off transcription or via multiple transcriptional terminators are significantly increased using divergent T7 RNAP promoters compared to convergent promoters. Finally, we have demonstrated that the spacer region between divergent promoters has no effect on the production of dsRNA when run-off transcription is used for termination, consistent with production utilising multiple transcriptional terminators in vitro.

### *In vitro* transcription using DNA templates with multiple transcriptional terminators reduces the formation of dsRNA multimers/aggregates compared to run-off transcription

IP-RP-HPLC analysis of dsRNA produced via IVT from DNA templates containing divergent promoters in conjunction with either transcriptional terminators or linearised at the end of the dsRNA gene resulted in differences in dsRNA product quality (see Figure 6). Differences in the peaks present with increased retention time after the main dsRNA product are observed for each of the dsRNAs. These peaks have previously been characterised as dsRNA multimers or aggregates (Nwokeoji et al. 2019). The results show a clear increase in the amount of the multimers and aggregates generated via run off IVT from linearised DNA templates compared to IVT from DNA templates containing multiple transcriptional terminators. Similar results were also obtained using convergent promoter systems (see Supplementary Figures S7).

**Figure 6.**
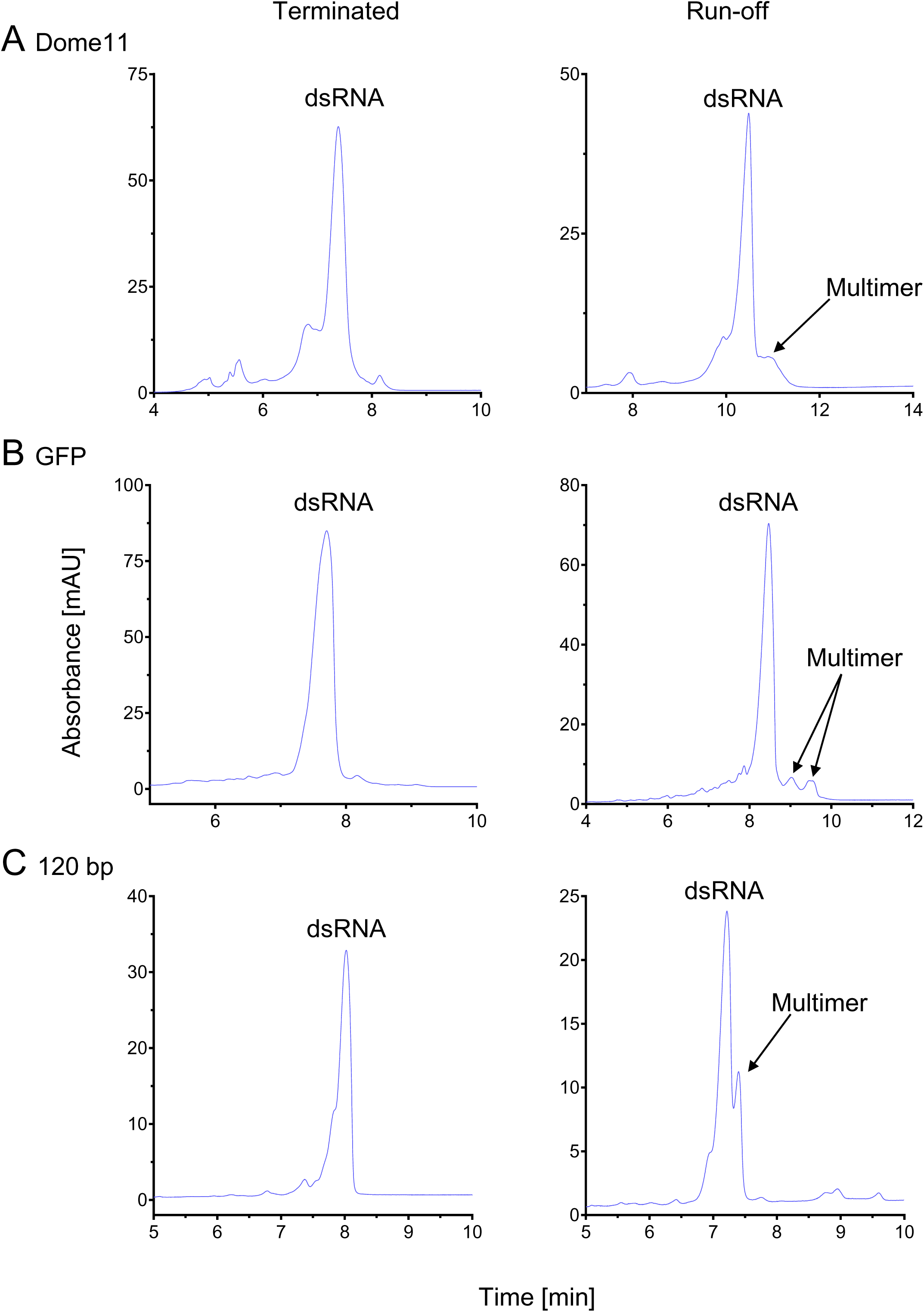
Comparative analysis of in vitro transcription of dsRNA produced from Divergent T7 polymerase DNA templates with and without multiple transcriptional terminators. IP-RP HPLC chromatography analysis of *in vitro* synthesised dsRNA of Dome11, GFP and 120 bp, utilising a divergent T7 polymerase. Samples were digested with DNase and RNase T1. Left chromatograms represent samples produced from terminated transcription. Right represents samples produced from run-off transcription. dsRNA main products and multimers are indicated. **A.** Dome11 divergent dsRNA samples. **B.** GFP divergent dsRNA samples **C.** 120 bp divergent dsRNA samples.

This is interesting to note, as the traditional production of mRNA and dsRNA by IVT is usually performed via run-off transcription (Dousis et al. 2023). Studies have demonstrated transcriptional extension by run-off transcription and the production of unintended dsRNA by-products (Piao et al. 2022; Dousis et al. 2023; Ziegenhals et al. 2023). It is proposed that 3’ extensions and resulting overhangs, may lead to increased disposition to form dsRNA multimers or even the formation of longer sequences. Further studies, such as denaturing IP-RP-HPLC, and atomic force microscopy, would allow us to confirm that these are indeed the formation of multimers, as previously performed by Nwokeoji et al. (2019).

However, previous data has demonstrated the production of multiple overhang products when producing dsRNA from IVT reactions using multiple transcriptional terminators without downstream digestion by RNase T1 (Ross et al. 2024). These findings demonstrate that higher quality dsRNA (reduction in dsRNA multimers/aggregates) is produced using multiple transcriptional terminators in comparison to run-off transcription only in conjunction with downstream processing with RNase T1.

## Conclusions

Double-stranded RNA plays an important role in a number of biological processes including gene silencing via the RNAi pathway. Furthermore, important therapeutic and agricultural applications of dsRNA are emerging, including siRNAs and dsRNA biocontrols. The successful deployment of dsRNA biocontrols requires efficient and cost-effective production of large quantities of dsRNA. Traditional production of dsRNA biocontrols is achieved *in vivo* in microbial cells or alternatively via large-scale in vitro transcription in conjunction with T7 convergent RNAP promoters. In this study, we designed a range of novel plasmid DNA constructs to study the effect of convergent and divergent T7 RNAP promoters for the production of dsRNA in microbial cells and via IVT.

*In vitro* transcription studies demonstrated that using divergent T7 RNAP promoters resulted in increased dsRNA yields compared to convergent T7 RNAP promoters for a wide range of different dsRNA sizes and sequences. It is proposed that the increased dsRNA yield using divergent T7 RNAP promoters is due to an elimination of T7 RNAP collisions when T7 RNAPs are transcribing in different directions, consistent with previous observations of bacterial and bacteriophage RNAPs (Callen et al. 2004; Crampton et al. 2006; Zhou and Martin 2006; Ma and McAllister 2009; Chatterjee et al. 2011; Hobson et al. 2012; Wang et al. 2023).

In addition, the results also showed that increasing the length of the spacer region (> 1kb) in the divergent T7 RNAP promoter system resulted in a significant decrease in dsRNA yield, potentially due to less efficient annealing of the ssRNA due to spatial proximity to each other. Furthermore, the results showed that the formation of dsRNA multimers was reduced when dsRNA was synthesised using IVT reactions, in conjugation with RNase T1, while using DNA templates with multiple transcriptional terminators, compared to run-off transcription. It is proposed that the decrease in dsRNA product quality is due to 3’ extensions following IVT, resulting in the formation of unintended dsRNA, or multimers.

The results from the production of dsRNA in microbial cells demonstrate that the optimal yields of larger dsRNA sequences are obtained using convergent T7 RNAP promoters. This would suggest that RNAP collisions do not have a major effect on the production of dsRNA *in vivo* (in contrast to *in vitro*). However, divergent T7 RNAP promoters were shown to be optimal for the smallest dsRNA sequence. It is proposed that smaller dsRNA sequences may experience more transcriptional interference due to a higher RNAP-to-DNA template base pair ratio during convergent T7 RNAP production. This potentially leads to increased collisions, which can terminate transcription and interfere with polymerase-promoter binding (Callen et al. 2004; Shearwin et al. 2005; Chatterjee et al. 2011; Wang et al. 2023).

It is important to note that *the in vivo* environment differs from the *in vitro* one. The effect of endogenous proteins, RNAs and other cellular machinery will play a role in transcription, annealing and degradation of target RNA species. Longer RNA sequences are more prone to degradation via nucleases and hydrolysis, as well as additional RNA protein binding sites that may hinder dsRNA annealing directly (Quendera et al. 2020; Wayment-Steele et al. 2021). Studies have also demonstrated that the stability of ssRNA, in the form of messenger RNA (mRNA), is negatively correlated with an increase in length (Chheda et al. 2024). These issues are potentially mitigated in the convergent production of larger dsRNA, due to the production of ssRNAs being localised nearer to one another on the plasmid within the cell.

Investigations into the spacer region between the divergent T7 RNAPs, showed that the size of the spacer region had no significant effect on dsRNA yields in microbial cells. Finally, these results highlight alternative optimum strategies for the production of a wide range of different sized dsRNA for therapeutic and agricultural applications.

## Materials and Methods

### Chemicals and reagents

Sigma Aldrich was used to source, ampicillin sodium salt, isopropyl β-D-1-thiogalactopyranoside (IPTG) ≥99%, LB Miller media, LB Broth with agar (Miller), sodium chloride (NaCl), sodium dodecyl sulphate (SDS) and tetracycline hydrochloride. Agarose gels were prepared with UltraPure agarose (Invitrogen) or molecular-grade agarose (Appleton). Agarose gel electrophoresis was performed using 1X Tris-acetate EDTA (TAE) buffer (Sigma Aldrich). Nucleic acid sample staining was prepared with Ethidium bromide (Alfa Aesar) or Midori green direct dye (Geneflow). RNA sample loading was aided with Novex™ TBE-Urea Sample Buffer (2X) (Thermo Fisher Scientific). Molecular cloning used the following kits: Genejet Plasmid Maxiprep kit (Thermo Fisher Scientific), GeneJet Gel extraction kit (Thermo Fisher Scientific), NEBuilder® HiFi DNA Assembly Master Mix (New England Biolabs), Quick Ligation™ Kit (New England Biolabs). Polymerase chain reactions were performed and purified using KAPA2G Fast Hotstart Readymix (Merck) and Monarch® PCR & DNA Cleanup Kit (New England Biolabs). Purification of DNA samples used UltraPure™ Phenol:Chloroform:Isoamyl Alcohol (25:24:1, v/v) (Thermo Fisher Scientific).

HiScribe™ T7 High Yield RNA Synthesis Kit (New England Biolabs) was used for *in vitro* transcription (IVT) reactions. N1-MethylpseudoUridine-5’-Triphosphate was sourced from BOC Sciences. IVT-produced RNA purification was performed with Monarch® RNA Cleanup Kit (New England Biolabs). DNA and ssRNA digestion was performed with TURBO DNase (Thermo Fisher Scientific) and RNase T1 (Thermo Scientific). HPLC mobile phases were prepared using UHPLC-MS grade acetonitrile (Thermo Scientific), UHPLC-MS grade water (Thermo Scientific), Triethylammonium acetate pH 7.4 (Sigma-Aldrich), ≥99.0% (GC) dibutylamine (Sigma-Aldrich), 99.5+% 1,1,1,3,3,3-hexaflouro-2-propanol (Thermo Scientific).

### Cell sources, growths and inductions

*E. coli* strains DH5α and HT115(DE3) were obtained from New England Biolabs and Jealott’s Hill International Research Centre, Syngenta, UK, respectively. The Mix and Go! *E. coli* Transformation Kit (Zymo Research) was used to generate competent cells, and transformations were performed following the manufacturer’s instructions. Plasmid preparation was performed using the Genejet Plasmid Maxiprep kit (Thermo Fisher Scientific), following an overnight growth of an individual colony in 250 ml of LB media. Growths and inductions of transformed HT115(DE3) cells to produce dsRNA were performed as previously described by Ross et al. (2024). Cell quantities were normalised to 1 x 10^9^ cells, calculated via the Agilent web tool, *E. coli* Cell Culture Concentration from OD_600_ Calculator (available at agilent.com/store/biocalculators/calcODBacterial.jsp).

### Polymerase chain reaction and DNA assembly

DNA fragments and synthetic genes were sourced from Integrated DNA Technologies and GeneArt® Gene Synthesis (Thermo Fisher Scientific), respectively. PCR reactions were performed with KAPA2G Fast Hotstart Readymix (Merck) and cycling parameters were optimised using the manufacturer’s guidelines. Overlapping PCR was performed via two sets of PCR amplification. The first set consisted of 15 cycles, with two fragments containing overlapping regions (16-32 bp). The second set (Extension PCR) consisted of 20 cycles, with the addition of two primers flanking the outer regions. Gibson assembly was performed using the NEBuilder® HiFi DNA Assembly Master Mix (New England Biolabs) according to the manufacturer’s instructions. Assembled products were then used as templates in standard PCR reactions prior to cloning. Purification of PCR products was performed with the Monarch® PCR & DNA Cleanup Kit (New England Biolabs) according to the manufacturer’s instructions.

### DNA template preparation and Cloning

DNA template preparation for IVT reactions and molecular cloning were performed with restriction enzymes sourced from Fisher Scientific and New England Biolabs. Cloning samples were purified with the GeneJet Gel extraction kit (Thermo Fisher Scientific) prior to ligation with the Quick Ligation™ Kit (New England Biolabs) and transformed as mentioned before. IVT templates were produced via plasmid linearisation with restriction enzymes or PCR, then purified with either the Monarch® PCR & DNA Cleanup Kit (New England Biolabs) according to the manufacturer’s instructions, Genejet Gel extraction kit or phenol/chloroform ethanol precipitation, as stated in Ross et al. (2024). Preliminary plasmid construct verification was performed via agarose gel electrophoresis as stated in Ross et al. (2024). Samples were sized using the DNA ladder GeneRuler 1 kb plus (Thermo Scientific). Sanger sequencing was used to verify further the plasmid constructs Genewiz (Azenta Life Sciences).

### RNA extraction and purification

RNA extractions and purifications were performed as described by Nwokeoji et al. (2016) with modifications previously stated in Ross et al. (2024). Additional modifications were made to the RNase T1 step for the removal of ssRNA during absolute quantification of dsRNA. A total of 2 μl of 1/10 diluted RNase T1 (1000 U/µL) (Thermo Scientific) was added to the clarified lysate and the incubation time was increased to 1 hour at 37 °C. Absolute dsRNA (total) was quantified and checked for contamination using a Nanodrop 2000c UV spectrophotometer (Thermo Fisher Scientific), with an absorbance factor of 46.52 μg/mL per A_260_ (Nwokeoji et al. 2017). Relative quantification was performed via weak IP-HPLC measuring the peak area (mAU*min) of the target full-length dsRNA unless stated otherwise. Quantitative analysis of gel band density was performed using the gel analyzer tool within the Fiji ImageJ software package (Schindelin et al. 2012).

### In vitro transcription

IVT reactions were performed using the HiScribe™ T7 High Yield RNA Synthesis Kit (New England Biolabs), following the manufacturer’s instructions. Equal molar quantities of template DNA (250 fmol), either from linearised plasmids or DNA fragments purified from gel extraction, were utilised in experiments. Samples were incubated for 2 hours at 37 °C. DNA templates were removed post IVT via the addition of 1 μL of TURBO DNase (2U/μL) (Thermo Fisher Scientific), incubated for 20 minutes at 37 °C. RNA products were purified via the Monarch® RNA Cleanup Kit (New England Biolabs), following manufacturers’ instructions, and eluted in 100 μL of RNase-free water. N1-MethylpseudoUridine-5’-Triphosphate was incorporated into reactions where mentioned, replacing Uridine at the same concentration stated in the HiScribe™ T7 High Yield RNA Synthesis Kit protocol.

### Ion pair-reverse phase high-performance liquid chromatography (IP-RP HPLC)

IP-RP HPLC analysis was performed on an UltiMate 3000 HPLC system (Thermo Fisher Scientific, UK, with a UV detection of 260nm, using a DNAPac RP column (2.1 x 10 mm or 2.1 x 100 mm, Thermo Fisher). Strong IP-RP-HPLC analysis was performed under the conditions: Buffer A, 15 mM dibutylamine (DBA) and 50 mM 1,1,1,3,3,3-hexaflouro-2-propanol (HFIP), Buffer B, 15 mM dibutylamine (DBA), 50 mM 1,1,1,3,3,3-hexaflouro-2-propanol (HFIP) and 50% acetonitrile (ACN). Strong IP-RP-HPLC gradients under native conditions (20°C) are as follows: Buffer B at 40% to 45% in 2 minutes, followed by a curved (4) extension to 65% for 14 minutes, up to 75% for 3 minutes at a flow rate of 0.17 ml/min. Buffer B at 35% to 40% in 2 minutes, followed by a curved (4) extension to 55% for 14 minutes, up to 75% for 3 minutes at a flow rate of 0.17 ml/min.

## Data availability

The data underlying this article are available in the article and in its online supplementary material or available on request.

## Supplementary data

Supplementary Data are available.

## Funding and Acknowledgements

SJR is a Biotechnology and Biological Science Research Council White Rose DTP iCASE student in collaboration with Syngenta (BB/T007222/1). MJD acknowledges further support from the Biotechnology and Biological Science Research Council (BB/M012166/1).

